# A Single-Cell Atlas of the Upper Respiratory Epithelium Reveals Heterogeneity in Cell Types and Patterning Strategies

**DOI:** 10.1101/2025.01.16.633456

**Authors:** Alexander G. Foote, Xin Sun

## Abstract

The upper respiratory tract, organized along the pharyngolaryngeal-to-tracheobronchial axis, is essential for homeostatic functions such as breathing and vocalization. The upper respiratory epithelium is frequently exposed to pollutants and pathogens, making this an area of first-line defense against respiratory injury and infection. The respiratory epithelium is composed of a rich array of specialized cell types, each with unique capabilities in immune defense and injury repair. However, the precise transcriptomic signature and spatial distribution of these cell populations, as well as potential cell subpopulations, have not been well defined. Here, using single cell RNAseq combined with spatial validation, we present a comprehensive atlas of the mouse upper respiratory epithelium. We systematically analyzed our rich RNAseq dataset of the upper respiratory epithelium to reveal 17 cell types, which we further organized into three spatially distinct compartments: the *Tmprss11a*+ pharyngolaryngeal, the *Nkx2-1*+ tracheobronchial, and the *Dmbt1*+ submucosal gland epithelium. We profiled/analyzed the pharyngolaryngeal epithelium, composed of stratified squamous epithelium, and identified distinct regional signatures, including a Keratin gene expression code. In profiling the tracheobronchial epithelium, which is composed of a pseudostratified epithelium-with the exception of the hillock structure-we identified that regional luminal cells, such as club cells and basal cells, show varying gradients of marker expression along the proximal-distal and/or dorsal-ventral axis. Lastly, our analysis of the submucosal gland epithelium, composed of an array of cell types, such as the unique myoepithelial cells, revealed the colorful diversity of between and within cell populations. Our single-cell atlas with spatial validation highlights the distinct transcriptional programs of the upper respiratory epithelium and serves as a valuable resource for future investigations to address how cells behave in homeostasis and pathogenesis.

**Highlights:**

- Defined three spatially distinct epithelial compartments, *Tmprss11a*+ pharyngolaryngeal, *Nkx2-1*+ tracheobronchial, and *Dmbt1*+ submucosal gland, comprising 17 total cell types
- Profiled Keratin gene expression code along proximal-distal and basal-luminal axes and highlighted “stress-induced” Keratins KRT6A and KRT17 at homeostasis
- Demarcated expression gradients of *Scgb1a1*+ and *Scgb3a2+* club cells along the proximal-distal axes
- Specified submucosal gland cell heterogeneity including *Nkx3-1+* mucin-producing cells, with ACTA2+ basal myoepithelial cells exhibiting gene profile for neuroimmune mediated signaling

## INTRODUCTION

The respiratory tract includes the nasal cavity, pharynx/larynx (throat), trachea, bronchi and lungs, functioning as a connected and interdependent unit. Mucosal surfaces lined by epithelial cells are essential elements of the respiratory tract, effective not only as a first-line physical barrier against chronic external threats, but also for host immune defense, and injury repair.^1–4^ The epithelium is vital for maintaining respiratory health. Conversely, it is also the site for pathogenesis of an array of upper airway acute and chronic conditions, such as laryngitis/pharyngitis, bronchitis, chronic cough, croup, etc.^5–7^ While substantial research has uncovered cellular diversity of the lower bronchopulmonary airways,^8, 9^ there is a considerable gap in knowledge on the upper respiratory epithelium, especially comprehensive analysis with regional comparisons. Many diseases demonstrate regional specificity in their manifestation, which underscores the importance of understanding the diverse transcriptomic profiles.

Research on the proximal conducting tracheal airway has identified significant cellular heterogeneity and revealed diverse roles in maintaining homeostasis^8, 10–12^ and facilitating repair.^3, 8, 11–13^ Current evidence suggests that the larynx plays a significant role in shaping host immunity and thus mediates respiratory health and disease.^14^ Yet, the laryngeal transcriptome remains relatively underexplored at the single cell level.^14^ Moreover, previous studies have predominantly focused on single-organ analyses, with no investigations to date that comprehensively examine both the trachea and larynx in tandem. While common cell types such as basal, ciliated, secretory, and club cells are found throughout both the larynx and trachea, their transcriptomic profiles have not been fully characterized to determine whether significant differences exist between these regions.

Submucosal glands (SMG) are primarily localized within the larynx, with a smaller reservoir situated in the ventral proximal trachea. These glands play a pivotal role in airway innate immunity by regulating the secretion of antimicrobial peptides and mucins essential for mucosal defense.^15–17^ Additionally, recent studies have identified rare, region-specific cell types with unique chemosensory and immune-mediated functions, including neuroendocrine cells,^18^ tuft cells,^19, 20^ and hillock structures.^21^ For example, tuft cells, known as brush cells in the trachea and solitary chemosensory cells in the nasal, pharyngeal, and laryngeal epithelium, exhibit dual roles in immune and chemosensory effector functions.^20, 22–25^ Neuroendocrine cells function as specialized sensory epithelial cells that protect the airways by releasing ATP to activate sensory neurons, triggering critical reflexes such as swallowing and expiration.^18^ Furthermore, rare tracheal hillock structures have been identified as injury-resistant reservoirs with distinct regenerative potential.^21^

With the variations in cell spatial patterning and the functional diversity of cell types, we hypothesized that the larynx and trachea possess unique, organ-specific transcriptomic profiles reflective of their specialized cell populations and tissue microenvironments. To analyze the cell diversity, we generated a comprehensive dataset covering the pharyngolaryngeal-to-tracheobronchial axis and established a single-cell atlas for the mouse adult upper respiratory epithelium. Our single-cell RNAseq (scRNAseq) analysis revealed 17 cell types across three spatially distinct compartments; (1) *Tmprss11a*+ stratified squamous epithelium (SSE) of the larynx, (2) *Nkx2-1*+ pseudostratified epithelium of the trachea, and (3) *Dmbt1+* SMG epithelium. Our dataset is rich with cell diversity, including abundant cell types, club (33%), basal (29%), and goblet-2 (11%), as well as rare cell types, including secretory-larynx (1%), tuft (0.5%), and neuroendocrine (0.1%) cells. We further separated basal cell transcriptomic signatures and gene-ontology analysis based upon respective compartment (trachea-dorsal:*Trp63+Cav1+),* (trachea-ventral:*Trp63+Tgm2+),* (larynx:*Trp63+Igfbp2+Tmprss11a+*), (SMG:*Trp63+Acta2+)*.

This analysis of basal cells, paired with regional profiling, highlighted a unique expression profile of Keratins and established 21 total expressed genes along the proximal-distal and basal-luminal axes. Most notably, previously identified “stress-induced” Keratins, KRT6A and KRT17, were found to be present during homoeostasis, with expression in all basal cells as well as luminal squamous cells. Additionally, our luminal cell analysis identified three distinct types of club cells, with higher levels of *Scgb1a1*+ and *Scgb3a2+* expression in more distal compared to proximal airway regions. Lastly, we profiled diversity of SMG cells, highlighting *Nkx3-1+* expression exclusive to goblet-1, goblet-2, and serous-acini cells, with ACTA2+ basal myoepithelial cells exhibiting a gene profile (*Ntng1, Ntn4, Slit3, Ntrk3, Cxcl12, Cxcl14*) linking potential neuroimmune strategies. Our single-cell transcriptomic analysis with *in situ* validations provides key insights into the transcriptomic and cellular heterogeneity of the upper airway and serves as a valuable atlas for hypothesis-driven work into responses to environmental insults, genetic mutations, and infectious diseases.

## RESULTS

### Single cell transcriptomic analysis of the upper respiratory epithelium reveal compartmental and cellular heterogeneity

To explore epithelial cellular diversity of the upper respiratory epithelium, we started by using scRNAseq to profile the pharyngolaryngeal-to-tracheobronchial axis. We isolated and pooled epithelial cells from pharynx to bronchi regions of adult wildtype C57BL/6 mice (n=4 males, 4 females), and applied them to 10x Chromium for scRNAseq (Fig. 1A). Following quality control (FigureS1A), 3,617lJepithelial single cells were recovered. Applying unsupervised clustering with Seurat default package,^26^ followed by supervised annotation using known markers (FigureS1A-D),^10, 13, 14, 17, 27^ we identified 17 cell types delineated with top markers (Fig. 1B; TableS1-3). These 17 cell types segregate into three compartments; pharyngolaryngeal, tracheobronchial, and SMG (Fig. 1C). In further analysis of the pharyngolaryngeal stratified squamous epithelium (SSE), we identified *Tmprss11a* as a novel marker for this compartment (Fig. 1H). The tracheobronchial pseudostratified epithelium was marked by *Nkx2-1,* a known marker that was region-specific, aside for its expression in the thyroid epithelium (Fig. 1I). In the SMG epithelia, we identified *Dmbt1* as a novel region-specific marker, which was expressed at a high level with negligible expression in a few club cells in the proximal trachea and subglottic surface epithelium (Fig. 1J). We validated these regional expression profiles using RNAscope on coronal sections along the pharyngolaryngeal-to-tracheobronchial axis, defining the spatial localization of the pharyngolaryngeal (*Tmprss11+),* tracheobronchial (*Nkx2-1)*, and SMG (*Dmbt1+)* compartments (Fig. 1D-J; FigureS1E). These expression patterns elucidate the transcriptomic specificity for regional targeting of these epithelial compartments.

**Fig. 1.**
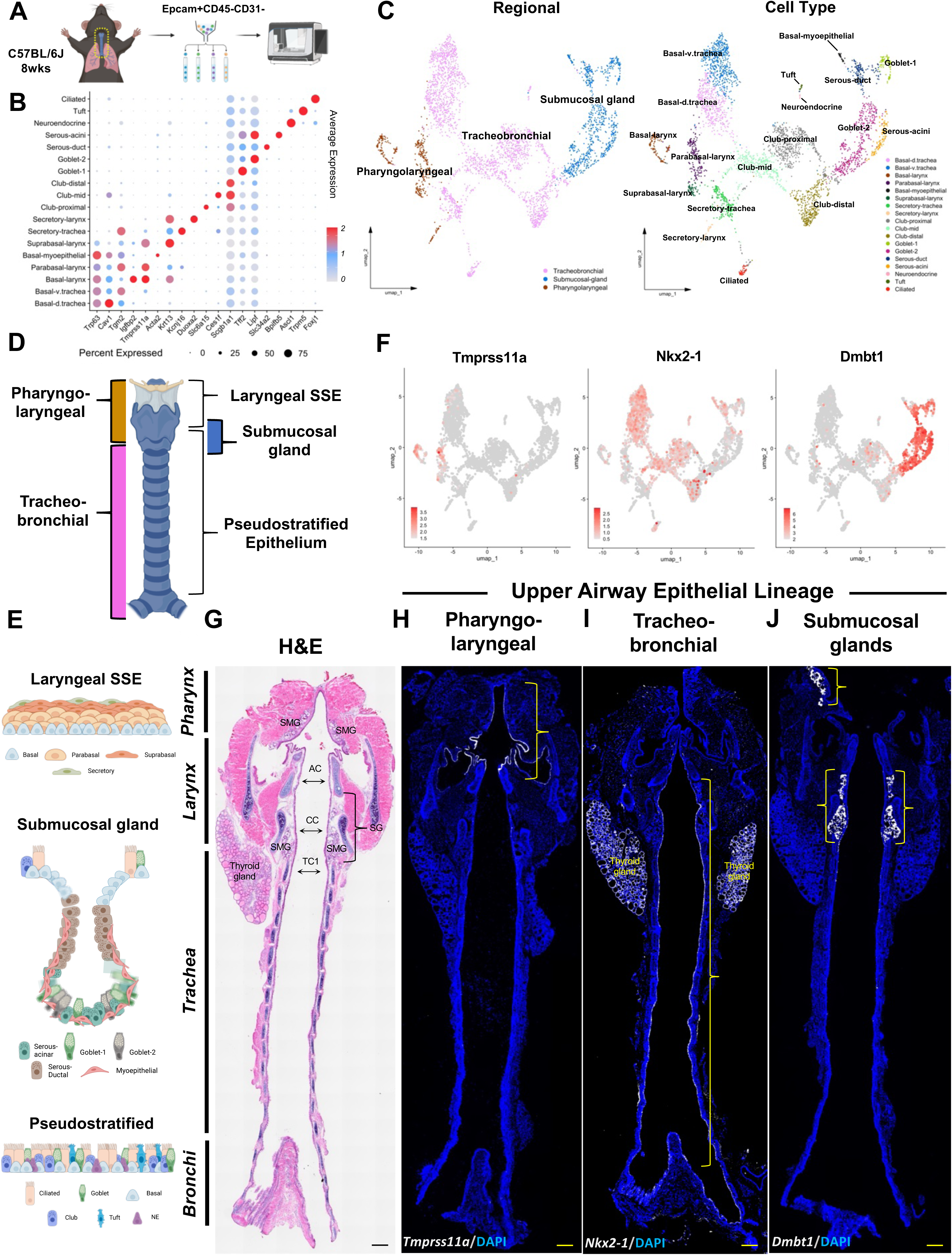
scRNAseq of the upper respiratory epithelium reveal compartmental and cellular heterogeneity. **(A)** Schematic illustration of cell isolation and scRNA-Seq workflow. **(B)** Dotplot showing top markers for each cluster. Using default Seurat^26^ dotplot settings (Methods), percentage expressed was plotted from 0 to 75% detected and the color bar shows the average of scaled normalized expression values across cells in a given cluster. **(C)** UMAP plots of integrated upper airway scRNA-Seq data from adult WT B6 mice (n=8 biological repeats, 4 male, 4 female). **(D,E)** Schematic illustration of macro-anatomical regions of the upper airway with compartmental epithelial cell type diversity. **(F)** Feature plots with **(H-J)** RNAscope validation of *Tmprss11a* (pharyngolaryngeal), *Nkx2-1* (tracheobronchial), *Dmbt1* (seromucous glands) exhibiting pan-epithelial regional markers in coronal sections**. (G)** H&E-stained upper airway. DAPI is in blue. All images 20X magnification. Scale bar represents 500μm. H&E hematoxylin and eosin, AC arytenoid cartilage, CC cricoid cartilage, TC tracheal cartilage, SSE stratified squamous epithelium, SG subglottis, SMG submucosal gland.

### Genome-wide profiling identifies signatures of the stratified squamous epithelium (SSE) along the basal-luminal axis

The pharynx and larynx are mostly lined with SSE, which caudally extends to the infraglottic vocal fold (VF),^28, 29^ and is also present in hillock regions of the trachea.^21^ This contrasts from the subglottis, trachea, and bronchi, which are lined with pseudostratified epithelium (Fig. 1D,E,G).^28, 29^ The multi-layered SSE serves to protect the underlying tissues from mechanical and chemical damage, pathogens, and dehydration.^21, 30^ Given the unique architecture, we hypothesized that this specialized SSE tissue would exhibit a basal-to-luminal coding pattern. Using SSE of the larynx as an example, we identified four cell clusters representing cell types along the basal-luminal axis: basal, parabasal, suprabasal/luminal and secretory (Fig. 2A,B). To explore the regional expression profile *in situ*, we utilized RNAscope and discovered that *Igfbp2* is exclusively expressed in the basal layer, *Tmprss11a* expressed in the basal and parabasal layers, and KRT13 protein distribution extending from basal to suprabasal with increased expression to differentiated luminal cells (Fig. 2B,C). We also identified populations of secretory cells dispersed throughout the pharyngolaryngeal apical epithelium, exhibiting top marker *Duoxa2* and *Il1a* expression (Fig. 1B; FigureS2C). Together these data define the compartmental specificity of squamous epithelia and unique expression profiles during terminal differentiation.

**Fig. 2.**
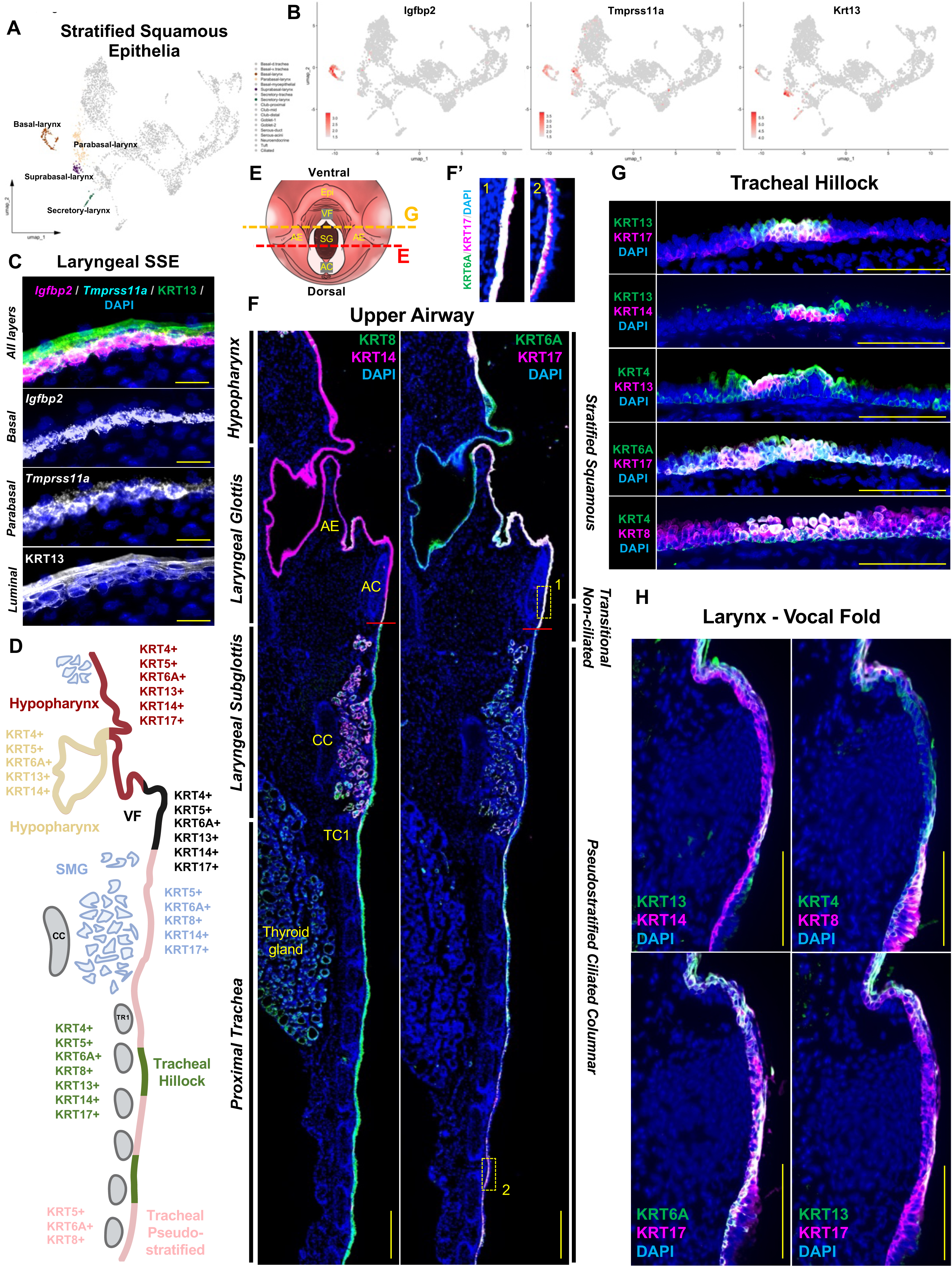
Genome-wide profiling revealed signatures of stratified squamous epithelium along basal-luminal axis with extensive Keratin diversity along proximal-distal axis. **(A)** Integrated UMAP plot highlighting laryngeal surface epithelial cells. **(B&C)** Feature plots with RNAscope validation of basal-to-luminal canonical markers for laryngeal SSE. **(D)** Schematic illustration of KRT diversity along the proximal-to-distal upper respiratory axis. **(E)** Schematic illustration of cranial view of larynx to display coronal sections (red/yellow dashed lines) attained for analysis. **(F,F’)** Immunofluorescent coronal serial sections with insets of dorsal upper airway epithelium displaying regional KRT expression along various epithelium subtypes. Red line indicates transition zone from SSE to pseudostratified. **(G,H)** Diversity of KRT expression to SSE of tracheal hillocks and laryngeal VF tissue. DAPI is in blue. All images 20X magnification. Scale bar represents 100μm (C,F,G) and 500μm (E). Epi epiglottis, AE aryepiglottic fold, SG subglottis, AC arytenoid cartilage, CC cricoid cartilage, TC tracheal cartilage, SSE stratified squamous epithelium, KRT keratin, SMG submucosal gland.

### Systematic profiling of Keratin code along the proximal-distal and basal-luminal axes

Keratins are essential for maintaining the structure and function of epithelial cells, playing pivotal roles in homeostasis, tissue repair, and disease pathogenesis.^31–39^ Specific Keratins are often markers for specialized epithelial cells, e.g. KRT13+KRT14+ for VF SSE and KRT4+KRT13+ for tracheal hillocks.^8, 21, 40^ However, the cell type-specific expression and spatial pattern of diverse Keratins have not been systematically characterized in the upper respiratory epithelium. We started with exploring Keratin gene expression in our scRNAseq dataset. Among the 63 known Keratin genes, expression of 21 of them were detected in the upper respiratory epithelium (FigureS3). Along the proximal-distal axis, we categorized the “Keratin code” based on scRNAseq data (Fig. 2D). We validated the data using immunofluorescence staining of key Keratins on serial sections (Fig. 2E-H; FigureS1F). The proximal-distal axis along the surface epithelium was defined by complementary KRT8 and KRT14 expression (Fig. 2F; FigureS1F).

Specifically, KRT8 was primarily detected in tracheal pseudostratified epithelium, SMG epithelium and thyroid gland epithelium, while KRT14 was detected primarily in laryngeal SSE, myoepithelial cells of SMG and a subset of basal cells of the proximal trachea (Fig. 2F-H; FigureS1F). KRT6A and KRT17, known as “stress-induced” Keratins,^31–33, 35^ were both highly expressed in all three compartments, defined by exclusive basal cell expression in pseudostratified and SMG epithelium, albeit, enriched in both basal and luminal cells in laryngeal SSE and tracheal hillocks (Fig. 2E-H; FigureS1F). Interestingly, and unique to the mouse, the lingual side of the aryepiglottic fold, which transitions ventrally into the epiglottis, contains cornified epithelium which did not express KRT17 (Fig. 2E,H; FigureS1F). We also compared the Keratin code in the VF and tracheal hillock squamous epithelium, two upper respiratory regions that are under constant mechanical insults. Keratin expression was abundant in both of these tissues with only subtle differences. Tracheal hillocks exhibited a Keratin coding pattern defined by basal markers (KRT4+KRT5+KRT6A+KRT14+KRT17+) and luminal markers (KRT4+KRT6A+KRT8+KRT13+) (Fig. 2F,H). VF SSE displayed identical basal marker Keratin expression to hillocks, and with identical luminal markers excluding KRT8 to the VF medial edge (Fig. 2F,G,H). Together, these data reveal a complex Keratin code in the upper respiratory epithelium (Fig. 2D; FigureS3).

### Basal cells exhibit transcriptomic regional diversity with unique functional signatures

Basal cells pave the entire upper respiratory epithelium, from pharynx to bronchi marked by high expression of the putative progenitor cell marker *Trp63* (Fig. 3A-C). To investigate the distinct patterning of these cells, we performed immunofluorescence and assessed *Trp63* expression in three compartments. Our analysis revealed multi-layered TRP63+ basal cells in the SSE of the larynx, single-layered basal cells in the trachea and subglottis and distributed basal cells in the SMG (Fig. 3B,B’). To further explore functional differences based on these anatomic locations, we isolated out basal populations *in silico* (Fig. 3D) and performed differential expression (DE) analysis (p_adj = <0.01, avg_logFC = 2.5), comparing across basal subtypes (Fig. 3E), followed by Gene Ontogeny (GO) and irGSEA hallmark pathways analysis (Fig. 3E-H; FigureS2A). The irGSEA hallmark pathway analysis, based on DE genes across all basal groups, found unique profiles within the basal-larynx, basal-myoepithelial, and basal-tracheal cell populations. Basal-larynx cells were enriched for G2M-Checkpoint, MYC-Targets-v1/v2, and DNA-Repair (*Krt13, Igfbp2, Top2a, Mki67*). Basal-myoepithelial cells were enriched for Epithelial-Mesenchymal-Transition and Apical-Surface (*Ntng1, Sox9, Acta2*, *Ntrk3*). Lastly, basal-tracheal cells showed enrichment for Inflammatory-Response and TNFα-Signaling-via-NF-κB (*Aqp5, Krt19, Scgb3a2, Cyp4b1, Lgr6*) (Fig. 3E,F). To rigorously assess these transcriptomic signatures, we conducted Gene Ontology (GO) enrichment and identified relative DE genes associated with KEGG pathway terms (Fig. 3G,H; FigureS2A; TableS4). Basal-larynx cells showed top enriched pathways related to Cell Cycle, p53 Signaling Pathway and Cellular Senescence (*Cdc20*, *Ccna2*, *Ccnb2*, *Ccnb1*, *Cdk1*, *Bub1b*). Basal-tracheal cells exhibited enrichment for Bile Secretion, N-glycan Biosynthesis, and Cortisol Synthesis and Secretion (*Nceh1*, *Aqp4*, *Atp1b1, Mgat4a, Mgat3, Pde8a*). Meanwhile, basal-myoepithelial cells showed enrichment for Focal Adhesion, Axon Guidance, and PI3K-Akt Signaling Pathway (*Ntng1*, *Epha4*, *Epha7*, *Sema6a*, *Cxcl12*, *Wnt5b*, *Ntn4*, *Slit3*, *Pak3*, *Myl9*) (Fig. 3G,H). For a comprehensive list of GO enrichment and irGSEA hallmark pathways analysis of DE genes across all epithelial cell types refer to TableS3 and FigureS4, respectively. This analysis highlights the distinct transcriptional and functional specialization of basal cell populations across the upper respiratory epithelium, providing a foundation for understanding their unique roles in homeostasis, repair, and regional adaptation.

**Fig. 3.**
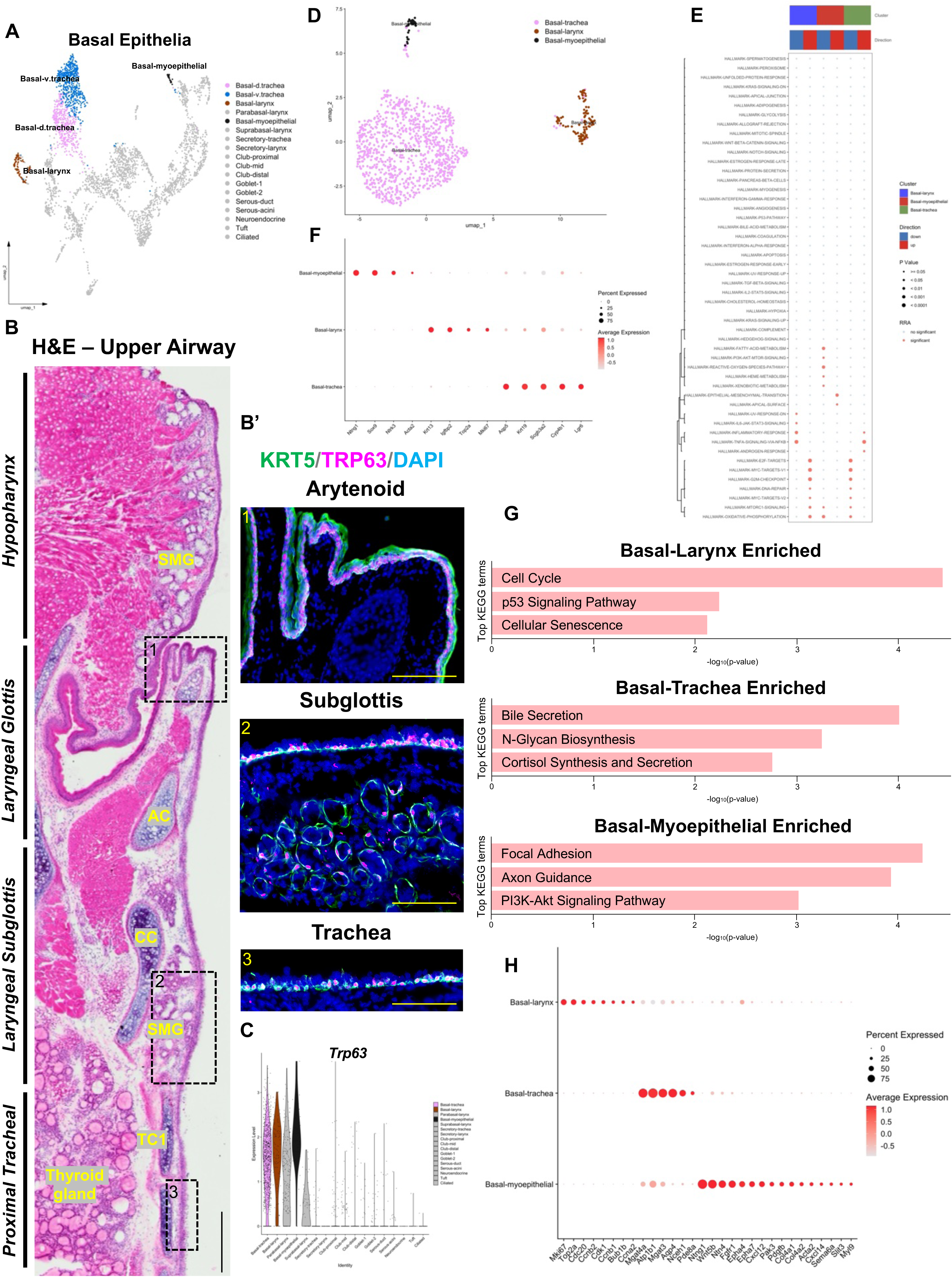
Basal cells exhibit transcriptomic regional diversity with unique functional signatures. **(A)** Integrated UMAP plot highlighting basal epithelial cells. **(B&B’)** H&E-stained upper airway with immunofluorescence exhibiting regional KRT5+TRP63+marked basal cells. **(C)** Violin plot exhibiting cell types that express canonical basal cell marker *Trp63*. **(D)** Integrated UMAP of basal cell subset. **(E)** irGSEA hallmark pathways analysis of DE genes across our three basal populations. **(F)** DotPlot showing top DE genes comparing basal-larynx (*Krt13, Igfbp2, Top2a* and *Mki67*) to basal-trachea (*Aqp5, Krt19, Scgb3a2, Cyp4b1* and *Lgr6*) to basal-myoepithelial (*Ntng1, Sox9, Ntrk3* and *Acta2*) in our basal cell subsetted scRNAseq dataset. **(G)** Gene Ontogeny (GO) enrichment analysis displaying top KEGG pathway terms enriched in each basal cell. **(H)** DotPlot showing top DE genes associated with functional enrichment analysis. DAPI is in blue. All images 20X magnification. Scale bar represents 100μm. H&E hematoxylin and eosin, KRT keratin, AC arytenoid cartilage, CC cricoid cartilage, TC tracheal cartilage, SMG submucosal gland.

Basal cells are progenitor cells that are critical for repair and regeneration of the epithelium following injury by differentiating into specialized cell types needed for tissue integrity. To compare the potential differentiation pathways in each of the basal cell types, we performed single cell trajectory analysis (Monocle 3). *In silico,* we selected cell clusters to only include each basal cell population (i.e. basal-larynx, basal-trachea, basal-myoepithelial) alongside the differentiated cell types in the corresponding region (FigureS5A-C). The basal-larynx trajectory revealed a direct lineage progression from basal to parabasal to suprabasal cells, followed by terminally differentiated cells of secretory, ciliated and club identities. The basal-trachea trajectory displayed a sequential progression through club-mid, club-proximal, and club-distal stages, followed by secretory lineages, with ciliated cells representing the most distal point in pseudotime. In comparison, basal-myoepithelial cells followed a SMG lineage trajectory progressing through goblet-2, serous-acini, serous-duct, goblet-1, ending in ciliated cells (FigureS5A-C). Our data highlights that basal cells, despite their morphological similarity, exhibit transcriptional heterogeneity primed for shared but also distinct roles across different epithelial compartments.

### Submucosal gland cell heterogeneity as revealed at single cell resolution

We sought to explore the heterogeneity of the murine SMG compartment, which has largely been unexplored at the single-cell level.^3^ Our scRNAseq revealed the expected rich diversity of differentiated cell types in the SMG, and also uncovered markers for each population (Fig. 4A,B). In the mouse upper respiratory tract, SMG were found in the pharynx, rostral laryngeal side of the epiglottis, subglottis, and proximal ventral trachea (Fig. 4C,D). Based on our scRNAseq data, these SMG epithelial cells segregate into multiple populations, including goblet-1 (*Muc5b, Tff2*), goblet-2 (*Lipf*), serous-duct (*Slc34a2*), serous-acini (*Bpifb5, Lipf*), and basal-myoepithelial cells (*Acta2, Trp63*), each with a distinct transcriptional profile (Fig. 4A,B). *Sox9* transcripts were also abundant in all SMG cells, with additional expression in the tuft cell population. Interestingly, RNAscope validation showed that the transcription factor *Nkx3-1* was expressed in goblet-1, goblet-2, and serous-acini cells (Fig. 4E), suggesting a unique role in regulating mucous producing cell fate. IllSMA+TRP63*+* basal-myoepithelial cells were found primarily in the distal acinar compartment and did not overlap with *Nkx3-1* expression (Fig. 4E). These data delineates both shared and distinct signatures of murine SMG cell types.

**Fig. 4.**
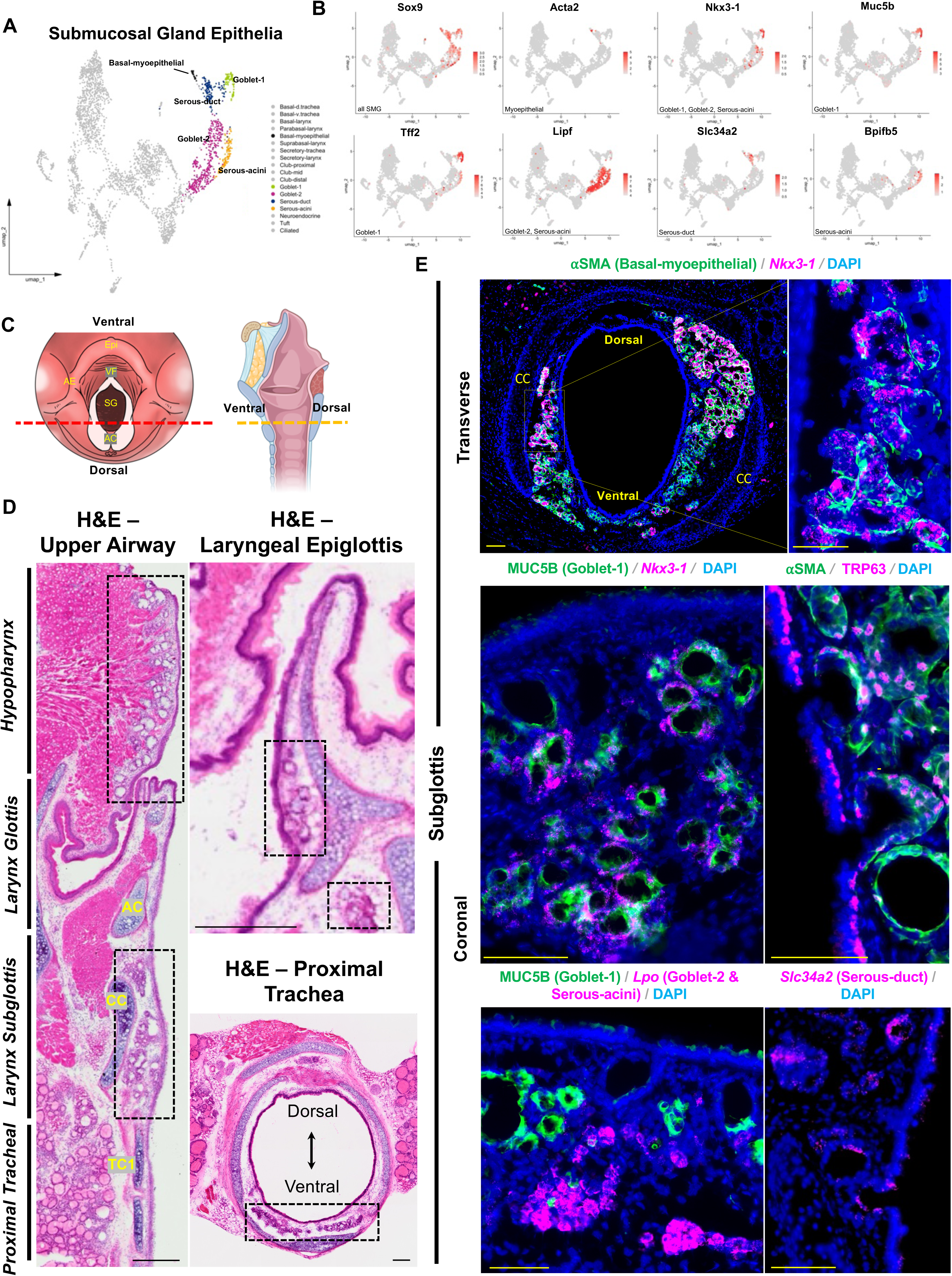
Submucosal gland cell heterogeneity as revealed at single cell resolution. **(A)** Integrated UMAP plot highlighting SMG epithelial cells. **(B)** Feature plots of canonical markers for SMG epithelial populations. **(C)** Schematic illustration of cranial and sagittal view of larynx to display coronal and transverse sections (red/yellow dashed lines) attained for analysis. **(D)** H&E-stained upper airway displaying SMG compartments (dashed black boxes). **(E)** Immunofluorescence transverse and coronal sections of SMGs displaying cell type canonical markers and unique transcription factor *Nkx3-1*. DAPI is in blue. All images 20X magnification. Scale bar represents 100μm. H&E hematoxylin and eosin. Epi epiglottis, AE aryepiglottic fold, SG subglottis, AC arytenoid cartilage, CC cricoid cartilage, TC tracheal cartilage.

### Luminal cell heterogeneity as revealed at single cell resolution

We further analyzed our scRNAseq dataset to reveal the diversity of luminal cell populations along the pharyngolaryngeal-to-tracheobronchial axis (Fig. 5A-C). Luminal cell types located to laryngeal SSE include suprabasal cells (KRT13, KRT17), secretory cells (*Duoxa2*, *Il1a*), and innervated neuroendocrine cells (*Ascl1,* SNAP25, GNAT3) within taste buds (Fig. 5C; FigureS2B,C). Solitary neuroendocrine cells were also found within the highly innervated subglottic epithelium and decreased in number along the proximal-distal axis towards the tracheal epithelium (Fig. 5C; FigureS2B). Luminal cell types located in the trachea epithelium include club (SCGB1A1, SCGB3A2), secretory/goblet (AGR2), ciliated (A-TUB), tuft (DCLK1, *Chat*), and hillocks (KRT13, KRT17). Ciliated cells were found throughout the pseudostratified epithelium of the trachea and in focal regions within the laryngeal epithelium, such as the superior VF and rostral lingual side of the epiglottis (not shown). We found heterogeneity within club cells along the tracheobronchial axis which we termed club-proximal, club-mid and club-distal, with subtypes showing unique regional transcriptomic signatures (Fig. 5A; FigureS2D-I).

**Fig. 5.**
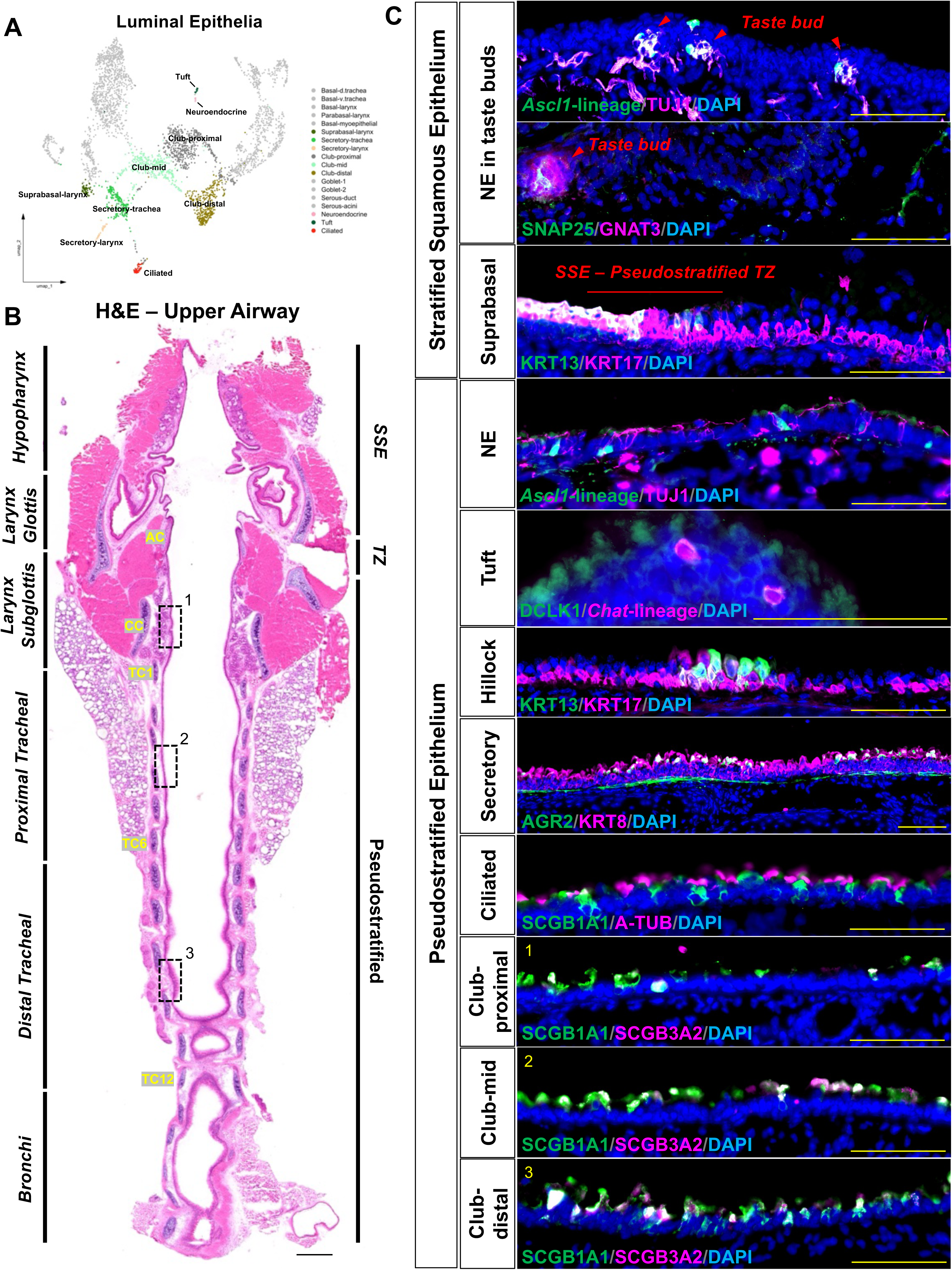
Luminal cell heterogeneity along pharyngolaryngeal-to-tracheobronchial axis revealed at single cell resolution. **(A)** Integrated UMAP plot highlighting luminal epithelial cells. **(B)** H&E-stained upper airway displaying anatomic regions of interest with dashed black boxes representing localized regions for club-proximal, club-mid, and club-distal immunofluorescent corresponding images. **(C)** Immunofluorescence images exhibiting unique markers for luminal cell types of the upper airway. DAPI is in blue. Images are either 20X or 40X magnification. Scale bar represents 100μm (C) and 500μm (B). H&E hematoxylin and eosin, AC arytenoid cartilage, CC cricoid cartilage, TC tracheal cartilage, TZ transition zone, NE neuroendocrine, SSE stratified squamous epithelium.

To directly compare DE genes, we took a similar approach as our basal analysis and isolated club cells *in silico*, and subclustered the population (FigureS2G; TableS5). Notably, distal club cells expressed the highest levels of *Scgb1a1* and *Scgb3a2* (Fig. 1C; FigureS2D,H), which was validated by pairwise DE comparison analysis (p_adj = <0.05, LogFC = 1) between club-proximal and club-distal (FigureS2F; TableS5). *Slc6a15* emerged as the top marker for club-proximal population across all cell types (Fig. 1C), which also showed elevated expression of *Cadm2* (FigureS2H; TableS1), a key defining marker expressed in laryngeal club cells.^14^ Immunofluorescence confirmed an increase in SCGB1A1+SCGB3A2+ club cell abundance to the distal trachea and extrapulmonary bronchi compared to proximal regions (Fig. 5C; FigureS2E). GO enrichment and pathway analysis revealed the top KEGG terms for club-proximal cells, including Ras Signaling Pathway (*Angpt1, Abl1, Gab2, Gnb5*), and Calcium Signaling Pathway (*Chrm3, Gna14, Erbb4, Tacr1*). In contrast, club-mid cells showed enrichment for KEGG terms related to Inflammatory Mediator Regulation of TRP Channels (*Camk2d, F2rl1, Calml3*) and Mucin-Type O-Glycan Biosynthesis (*Galntl6, B4galt5*). Finally, club-distal cells were enriched for KEGG terms linked to Ribosome function and Coronavirus Disease (*Mrpl4, Rps26, Mrpl2, Rps19, Rps7, Rpl36al, Rplp0, Rps12*) (FigureS2I; TableS5). Together, our analysis suggests potential regional-specific roles for club cells, where club-proximal cells located in the highly innervated laryngeal subglottis may play a prominent role in xenobiotic detection, while club-mid and club-distal cells localized to the trachea are primarily involved in secretory/mucin pathways and metabolic processes.

## DISCUSSION

To date, research has established diverse heterogeneity across the cellular landscape of the unified respiratory epithelium,^8, 9, 14^ offering an initial basis for its complexity. Building on these investigations focused on individual organs, here we present a comprehensive analysis of the single cell landscape in the murine upper airway, encompassing the pharyngolaryngeal-to-tracheobronchial axis. We defined three spatially distinct tissue compartments *Tmprss11a*+ pharyngolaryngeal, *Nkx2-1*+ tracheobronchial, and *Dmbt1*+ SMG epithelium, comprised of 17 transcriptionally unique cell types. This comprehensive regional atlas provides a critical framework for future studies aimed at understanding cellular dynamics in airway health, disease progression, and repair mechanisms.

Our analysis revealed a robust population of TRP63+ basal cells, characterized by regional transcriptomic diversity and distinct functional signatures. Consistent with prior studies demonstrating basal cell subpopulations in dorsal versus ventral tracheal regions,^13^ we identified a dorsal-enriched population (*Trp63+Cav1+*) and a ventral-enriched population (*Trp63+Tgm2+*) of tracheal progenitors. In the larynx, basal cells exhibited unique enrichment for *Igfbp2*, with a concomitant reduction in *Tmprss11a* expression among SSE progenies.

Laryngeal SSE display higher cellular turnover compared to the predominantly quiescent pseudostratified epithelium.^21, 41^ Comparative analyses between basal cells from the larynx and trachea revealed elevated expression of proliferative markers in laryngeal basal cells suggestive of key roles in tissue repair, while tracheal basal cells enriched for markers associated with secretory function indicate their contribution to mucosal defense. These findings highlight distinct regional specialization of basal cell populations between the larynx and trachea, underscoring their functional divergence in maintaining epithelial integrity.

We also explored Keratin gene expression within our scRNAseq dataset with the goal of establishing a comprehensive Keratin profile of the upper respiratory epithelium. Systematic profiling identified a total of 21 represented Keratin genes, delineating a distinct expression code along both the proximal-distal and basal-luminal axes. Among these, we focused on the “stress-induced” Keratins, KRT6A and KRT17. While previous studies have reported ectopic expression of KRT6A and KRT17 in epithelium following wound healing^35^ or in the context of viral persistence,^32, 33^ our findings reveal widespread expression under homeostatic conditions. Specifically, KRT6A+KRT17+ epithelium was detected in basal cells of the trachea and SMG, as well as in both basal and luminal cells of the laryngopharynx and tracheal hillocks. We also observed restricted expression of KRT8 to pseudostratified surface epithelium and KRT14 to squamous surface epithelium of the pharyngolarynx, delineating a distinct proximal-distal axis of epithelial differentiation. KRT13 at the protein and gene level marked suprabasal (i.e. luminal) SSE, however, was also modestly expressed in our basal-larynx population. Recent work has reported KRT13 expression in basal cells of hillocks.^21^ Immunofluorescent validation found hillocks expressed a near identical Keratin code as VF SSE, thus we suspect that hillocks share a sufficiently similar transcriptome with differentiated SSE, leading them to be grouped into our suprabasal-larynx population. To date, no studies have investigated airway SSE of both the larynx and trachea within one dataset at single cell resolution. These findings highlight previously uncharacterized Keratin dynamics across epithelial subtypes, providing new insights into airway squamous epithelial differentiation and potential shared transcriptional profiles with specialized structures like hillocks.

Next, we sought to characterize the cellular diversity of an underexplored, yet essential tissue compartment, the SMG. SMG cells play critical roles in upper airway immune defense, actively secreting mucus and antimicrobial byproducts to defend against pathogens and irritants, and with unique basal-myoepithelial cells.^16, 42^ We identified *Nkx3-1* transcription factor, uniquely expressed in goblet-1, goblet-2, and serous-acini cells. *Nkx3-1*, widely recognized as a prostate epithelium-specific marker, plays a key role in the development of the testes and prostate and has been implicated in prostate cancer.^43, 44^ Given its cell-specific expression and well-established roles in development, we hypothesize that this transcription factor may influence the cell fate of mucin-producing cells in the SMG, potentially serving as an effective candidate for disease modeling. The subglottic epithelium shares a close anatomical and functional relationship with adjacent SMG. Notably, *Acta2+* myoepithelial cells within SMG function as reserve stem cells capable of regenerating the airway surface epithelium.^3, 45, 46^ In salivary glands, myoepithelial cells have been identified as key sources of neurotrophin signaling following irradiation-induced damage.^47^ When comparing basal-myoepithelial cells from the larynx and trachea, we observed increased expression of genes associated with axon guidance (*Ntng1, Wnt5b, Ntn4*), cytokine signaling (*Cxcl12, Cxcl14*), and neurotrophic receptors (*Ntrk2, Ntrk3*). The *Cxcl12-Cxcr4* axis is a well-characterized signaling pathway with pleiotropic roles in immune responses, viral infections, and cancer.^48, 49^ Recent studies have further demonstrated that *Cxcr4* is upregulated following influenza infection, modulating neutrophil phagocytic activity, ligand-specific migration, and the induction of neutrophil extracellular traps.^50^ Based on our findings, we propose that basal-myoepithelial cells may play a critical role in injury repair and immune response mediated by the *Cxcl12/14-Cxcr4* axis. In addition, we detected upregulation of CHAT muscarinic receptors (*Chrm1, Chrm3*) in basal-myoepithelial cells, suggesting potential crosstalk with cholinergic-expressing neurons in the airways. These findings highlight the intimate relationship between basal-myoepithelial cells and innervating neurons and suggest a critical role for these cells in mediating neuroimmune interactions that contribute to airway homeostasis and disease.

The upper respiratory epithelium of the larynx and trachea contains several shared luminal cell types, including club, secretory, and ciliated cells. However, the transcriptomic differences between these regions remain incompletely characterized. To address this, we hypothesized that distinct transcriptomic profiles would emerge based on anatomical localization. Our analysis identified three transcriptionally distinct populations of club cells, with higher levels of *Scgb1a1*+ and *Scgb3a2+* expression in more distal compared to proximal regions along the upper airway axis. Club cells, defined by secretoglobin family genes (*Scgb1a1, Scgb3a1, Scgb3a2*), have been shown to exhibit functional heterogeneity in previous studies.^8, 51, 52^ A unique club-proximal population, exclusively expressing the defining laryngeal club cell marker *Cadm2*, was localized to the larynx.^14^ Additional populations, designated as club-mid and club-distal, were restricted to the proximal trachea and distal trachea/extrapulmonary bronchi, respectively. Notably, laryngeal club cells, which have only recently been described, exhibit both squamous and cuboidal morphologies,^28^ and are implicated in detoxification, repair, regeneration, and anti-inflammatory processes.^14, 52, 53^ We also identified a novel, rare secretory cell population within the larynx, marked by high expression of *Duoxa2*, an enzyme critical for thyroid hormone synthesis and mucosal immunity.^54^ This rare cell population exhibited additional upregulated genes, including mucins (*Muc13*), interleukin signaling components (*Il1a, Il1r2, Il1rn*), the transcription factor *Irf7*, and genes associated with inflammatory responses and pathogen defense (*Isg15, Tnf, Tnfrsf11b*). The unique transcriptional profile of this cell type suggests a potential role in coordinating mucosal defense and immune responses against pathogens in the laryngeal epithelium.

In addition to commonly shared cell types, several rare yet functionally significant populations exist in the upper airway, including neuroendocrine cells, tuft cells, and ionocytes, each contributing uniquely to airway homeostasis and function. Our dataset identified both neuroendocrine and tuft cells; however, ionocytes were not detected, likely due to their scarcity and potential methodological differences in tissue dissociation compared to prior studies.^8^ Solitary neuroendocrine cells expressed canonical markers (*Ascl1, Calca*)^8, 55^ and exhibited elevated expression of *Cxcl13* and *Ngf*, with the highest abundance within the laryngeal subglottic epithelium. Neuroendocrine cells in the lung, known as specialized airway sensors, are critical for interacting with peripheral nerves to regulate immune responses to allergens.^56, 57^ Tuft cells in our dataset were marked by classical tuft cell genes (*Pou2f3, Spib, Trpm5, Gnat3, Dclk1*) along with genes associated with immune regulation (*Rgs13, Rgs14, Rgs21, Tnfrsf13b, Il17rb*)^14^ and basal cell identity (*Lgr5*). Although limited cell numbers precluded identification of subpopulations, other studies have reported heterogeneity within tuft cells, including tuft-1 (*Pou2f3*) and tuft-2 (*Pou2f3, Gfi1b, Spib, Sox9*) subtypes in the tracheal epithelium.^8, 22^ Tuft cells have also been characterized as sources of IL-25, prostaglandins, leukotrienes, and acetylcholine,^20, 23–25, 58, 59^ which are crucial for promoting innate type 2 immunity and protective neural reflexes.^60^ Our use of Chat^cre^;tdTomato lineage-marked tissue confirmed CHAT expression in tracheal tuft cells, supporting their acetylcholine-producing capacity and aligning with previous work.^61, 62^ These findings underscore the extensive heterogeneity of epithelial cells across multiple upper airway compartments, emphasizing their roles in defending against environmental insults and maintaining homeostasis.

The extensive transcriptomic and cell diversity of the upper respiratory tract enables the epithelium to perform a range of functions, from forming physical barriers and initiating immune responses to repairing tissue damage and regulating inflammation. Our findings establish, for the first time, a region-specific specialization of epithelial diversity and a detailed cross-organ single-cell atlas along the pharyngolaryngeal-to-tracheobronchial axis, offering valuable insights into the diverse cell populations that maintain airway health and resilience.

## METHODS

### Animals

All experimental procedures were performed in the American Association for Accreditation of Laboratory Animal Care (AAALAC)-certified laboratory animal facility at the University of California, San Diego (UCSD). All animal husbandry and experiments were conducted under approved Institutional Animal Care and Use Committee (IACUC) guidelines. Wild-type C57BL/6 (JAX 000664)*, Ascl1^CreERT^*^2^ (JAX 012882), *Rosa^lxl-tdTomato^*(*Ai14*, JAX 007914), and *Chat^cre^* (JAX 031661) lines were purchased from the Jackson lab. All the *cre* lines we used in this study were kept in C57BL/6 background. All *cre* driver lines in heterozygous form are viable and fertile, and no abnormal phenotypes were detected. Both male and female mice were used in the experiment. Adult mice were 8-10 weeks of age for all experiments.

### Tissue collection and immunofluorescence staining

Mice were euthanized by CO2 inhalation followed by transcardial perfusion with PBS to remove circulating blood. The larynx-trachea was isolated and fixed overnight in 1%PFA or immediately embedded in OCT, flash frozen using 2-Methylbutane and liquid nitrogen and stored at –80°C. Using a cryostat, the larynx-trachea was then sectioned (12 μm) and stored at –20°C. Unfixed tissue was then washed in PBS for 5mins to remove OCT, heated to boiling in 10 mM citrate buffer (pH 9) for antigen retrieval and treated with 0.5% Triton X-100 in PBS for 15mins. All sections were processed for immunostaining following a standard protocol.^28^ *Ascl1* recombination was induced via Tamoxifen IP administration with dose of 100mg/kg for three consecutive days. All primary and secondary antibodies used are listed in TableS6. Primary antibodies were applied overnight at 4°C, while secondary antibodies were applied for 1hr at room temperature. Sections were incubated with DAPI (1:1000 ratio) for 10 min at RT. Slides were mounted and coverslipped with Prolong Diamond mounting media (Fisher P36970), cured flat at room temperature in the dark for 24 h, and stored at 4°C. Each experiment was replicated at least twice for all timepoints and targets assessed.

### RNAscope *in situ* hybridization

All staining procedures were performed using the RNAscope Fluorescent Multiplex Kit V2 (Advanced Cell Diagnostics, no. 323100) following the manufacturer’s instructions. The following probes from Advanced Cell Diagnostics were used: Mm-*Tmprss11a* (no. 1232761), Mm-*Nkx2.1* (no. 434721), Mm-*Nkx3.1* (no. 472111), Mm-*Dmbt1* (no. 418561), Mm-*Igfbp2* (no. 405951), Mm-*Lpo* (no. 403521), Mm-*Slc34a2* (no. 449311), Mm-*Il1a* (no. 440398).

### Tissue processing and flow cytometry

Whole upper airway (larynx-trachea) was mechanically dissociated by mincing tissue with razor blades in solution containing 5Lml of RPMIL1640 (Thermo Scientific) with 10% fetal bovine serum, 1 mM HEPES (Life Technology), 1LmM MgCl_2_ (Life Technology), 1LmM CaCl_2_ (Sigma), 0.5LmgLml^−1^ collagenase D and type I/dispase (Roche), and 0.25Lmg DNaseLI (Roche). Minced tissue was then digested by shaking at around 150 rpm for 30 min at 37 °C. Following incubation, upper airway pieces were mechanically dissociated further by straining through a MACS 70 μm filter. Red blood cells were removed by the addition of 1 ml of RBC lysis buffer (BioLegend) to each tube and incubation at room temperature for 1 min. Single-cell suspensions were pelleted (1,500 rpm, 4 °C, 5 min), counted with a hemocytometer and diluted to around 1L×L10^6^ cellsLml^−1^. Diluted cells were stained with Fc blocking antibody (5LmgLml^−1^, BD). Cells were then incubated with specific surface marker antibody at 1:1000 APC-FITC-conjugated anti-Epcam (BioLegend, no. 118214) to isolate epithelial lineage. Cells were then stained using live/dead dye (CD11b: Brilliant Violet 450, BioLegend, cat#75-0112-U025) before being resuspended in 2% FBS + 1:2000 DAPI. All FACS sorting was done on a BD FACSAria Fusion Sorter (BD Biosciences) analyzer with three lasers (405, 488 and 640 nm) at the Flow Cytometry Core at VA San Diego Health Care System and San Diego Veterans Medical Research Foundation. All data were further analyzed and plotted with FlowJo software (Tree Star).

### 10x Chromium scRNAseq and data analysis

Single cells were processed into complementary DNA libraries using Chromium Single Cell 3’Lv3 kit (10X Genomics, Pleasanton, CA) and sequencing was carried out on the NovaSeq (Illumina) platform. The CellRanger software package from 10X Genomics (v3.0.2) was used to align raw reads onto the mouse reference genome (GRCm38) and generate single-cell gene barcode matrices. CellBender (v0.3.0)^63^ was then used to remove technical artefacts and ambient RNA to produce improved estimates of gene expression. The RLpackage Seurat (v4.0)^26^ was then used to perform data quality control, normalization, principal components analysis, UMAP generation and differential gene expression testing. To ensure the exclusion of potential thyroid epithelial contamination, we manually removed the thyroid gland during sample preparation and computationally filtered clusters expressing thyroid-specific genes (*Folr1, Tg, Foxe1*) from our dataset. Single cells with above 15% mitochondrial reads and between 200 and 7500Lunique genes were considered high-quality cells and were filtered for further analyses.

Additional filtering included a minimum UMI cutoff of 2 and minimum cell count cutoff of 10. LogNormalize global-scaling normalization method was applied with regressing variables cell cycle, S.Score, G2M.Score, and mitochondrial RNA level to remove confounding effect on downstream clustering. In addition, DoubletFinder (v2.0)^64^ was used to remove doublets and LogNormalize was again used to normalize feature expression. CCA^65^ was used to integrate dataset across two experimental batches for a total of n=8 mice (4 males, 4 females). In total 3,617Lepithelial single cells were recovered for analysis. To determine dimensions for optimized clustering, we evaluated the optimal cutoff using an elbow plot and settled on using the first 50Lprincipal components for clustering and projection with UMAP. Clustering resolutions was set to 0.8 (Seurat default), resulting in 19Linterim clusters representing potential unique cell populations. Following manual inspection of top markers, we combined several clusters with shared markers to ensure that annotated clusters would show unique transcriptional profiles, resulting in 17Ldistinct clusters. For rigorous definition of marker genes for each cluster, we performed unsupervised and supervised analysis to screen each cluster’s top marker genes (Seurat, FindAllMarkers) using ViolinPlot, FeaturePlot and DotPlot. Cell clusters were referenced against available literature.^8, 14, 17^ Cell clusters were validated with RNAscope and immunofluorescence staining (see main figures). We provide a list of the top 100Lmarker genes of all cell clusters in TableS1. High-throughput sequence data is available at GEO. Once exclusively differentially expressed genes were identified, we performed tests of enrichment using Gene Ontology (GO) annotations utilizing Enrichr (v2024)^66^ as previously described.^67^

---

**Figure S1. (A)** Quality control data including nFeature RNA (the number of genes detected in each cell), nCount RNA (the total number of molecules detected within a cell) and mitochondrial percentage of our integrated airway scRNAseq dataset. **(B)** Unsupervised clustering with Seurat default package for cell type annotation. **(C)** Supervised clustering exhibiting top known gene markers for cell type annotation. **(D)** Dendrogram displaying hierarchical relationships between clusters of cells using ‘PlotClusterTree’ Seurat package. **(E)** Top differentially expressed gene markers establishing macro-anatomical regional specificity. **(F)** Immunofluorescence of upper airway coronal serial sections in split-channel view indicating Keratins differential regional expression. DAPI is in blue. Images are 20X magnification. Scale bar represents 500μm. AE aryepiglottis, AC arytenoid cartilage, CC cricoid cartilage, SMG submucosal gland.

**Figure S2. (A)** Volcano plot exhibiting differential gene (DE) expression of basal cell populations. **(B)** 20X coronal immunofluorescent section from Ascl1^creERi/+^;ROSA^tdTom/+^ reporter murine tissue exhibiting *Ascl1*+lineage marked neuroendocrine cells (red) of the upper airway. **(C)** Secretory cells in pharyngolaryngeal apical epithelium exhibiting *Il1a* expression. **(D)** Feature plot exhibiting cell specific expression of *Scgb1a1* and *Scgb3a2* transcripts. **(E)** Immunofluorescence exhibiting increased abundance and expression of SCGB1A1+ and SCGB3A2+ proteins in distal and extrapulmonary bronchi. **(F)** Pairwise DE analysis of club-proximal versus club-distal exhibiting increased *Scgb1a1* and *Scgb3a2* expression to distal airway club cells. **(G)** Integrated UMAP of club cell subset. **(H)** Feature plot exhibiting cell specific expression *Scgb1a1*, *Scgb3a2, Scgb3a1,* and *Cadm2* transcripts from our club cell subsetted scRNAseq dataset. **(I)** DotPlot showing top DE genes associated with functional enrichment analysis. DAPI is in blue. Images are 20X magnification. Scale bar represents 100μm (C,E) and 500μm (B). AC arytenoid cartilage, CC cricoid cartilage, TC tracheal cartilage.

**Figure S3.** DotPlot exhibiting comprehensive profiling of Keratin diversity across all epithelial cell types in our integrated scRNAseq dataset.

**Figure S4.** irGSEA hallmark pathways analysis of differentially expressed (DE) genes across all epithelial cell types in our integrated scRNAseq dataset.

**Figure S5.** Subset UMAP plots of integrated scRNAseq dataset from upper airway analyzed with Monocle 3 trajectory mapping with pseudotime boxplot displaying **(A)** basal-larynx, **(B)** basal-trachea, and **(C)** basal-myoepithelial cell origins.

**Table S1.** Consensus marker genes for cell-type clusters from 3’ droplet-based scRNA-seq data (3,617 cells). FDR-corrected Fisher’s combined p-value < 0.001; Minimum avg_log2FC: 2.0; top n=100 genes.

**Table S2.** Cell-type enriched TFs, GPCRs and KRTs from 3’ droplet-based scRNA-seq data (3,617 cells). FDR-corrected Fisher’s combined p-value < 0.001; top n=500 genes.

**Table S3.** Functional enrichment analysis using upregulated genes of epithelial cells showing the top 10 biological processes, cellular components, and molecular functions.

**Table S4.** Table of significantly differentially expressed marker genes comparing basal-larynx versus basal-trachea versus basal-myoepithelial populations. Marker genes for each cell type from “FindAllMarkers” analysis are sorted by adjusted p-value, following the default Seurat pipeline and based on Bonferroni correction using all features in the dataset. KEGG pathway and GO Biological Process analyses (Enrichr (v2024)) exhibiting Top 10 upregulated terms in each group.

**Table S5.** Table of significantly differentially expressed marker genes comparing club-proximal versus club-distal populations. Marker genes for each cell type from “FindAllMarkers” analysis are sorted by adjusted p-value, following the default Seurat pipeline and based on Bonferroni correction using all features in the dataset. KEGG pathway and GO Biological Process analyses (Enrichr (v2024)) exhibiting Top 10 upregulated terms in each group.

**Table S6.** Antibodies and reagents used in this study.

## Acknowledgements

We thank J. Sabatini and M. Erb (UCSD School of Medicine Microscopy Core, supported by NIH grant NINDS P30NS047101) for help with scanning microscopy. This work was supported by grants NIH NHLBI R01 AT011676-01, 1R01 HL160019-01, and NIH NIDCD F32 DC021634-01.

## Author contributions

Conceptualization: A.G.F, X.S. Methodology: A.G.F. Investigation: A.G.F. Visualization: A.G.F Funding acquisition: A.G.F., X.S. Supervision: X.S. Writing – original draft: A.G.F. Writing – review & editing: A.G.F., X.S.

## Declaration of interests

The authors declare no competing interests.

## Inclusion and diversity

We support inclusive, diverse, and equitable conduct of research.

## Data and code availability

All data generated supporting the findings of this study are available in the manuscript. Further information is available from the lead author upon reasonable request.

This paper does not report original code.

## Notes

### Competing Interest Statement

The authors have declared no competing interest.

