## Supplementary figures and images for "A Single-Cell Atlas of the Upper Respiratory Epithelium Reveals Heterogeneity in Cell Types and Patterning Strategies"

### Supplemental figures

Foote, Figure S1

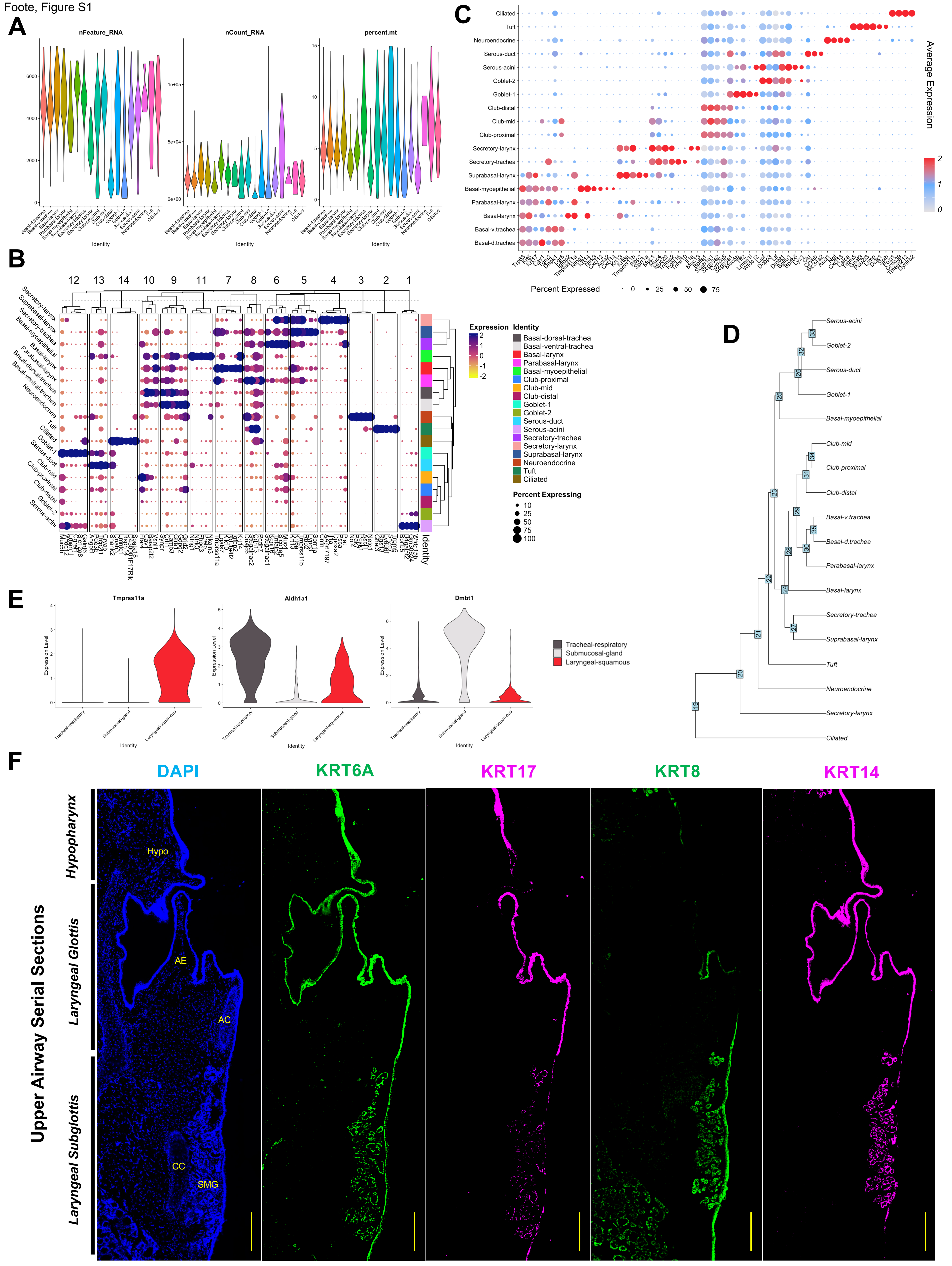

Foote, Figure S2

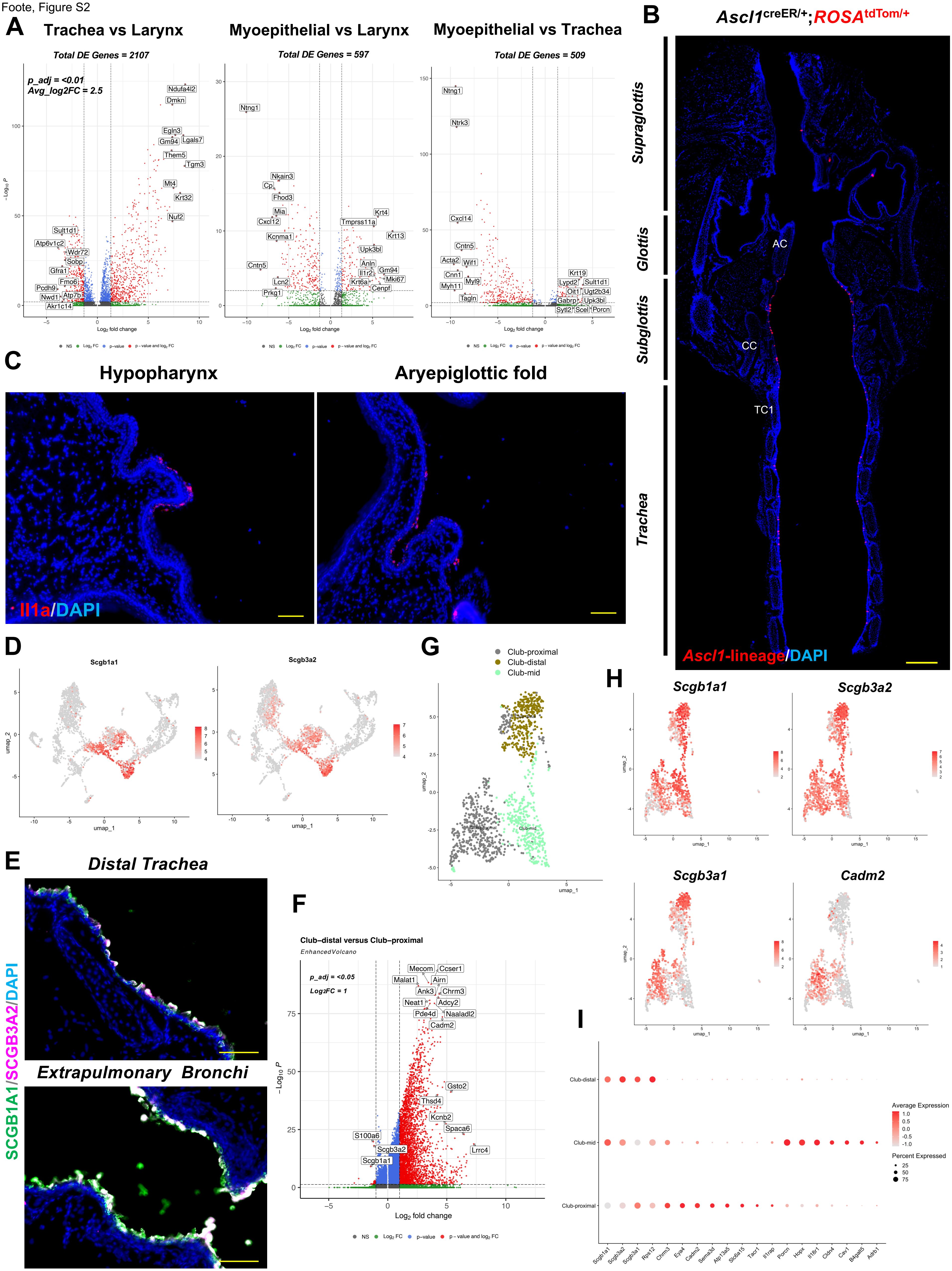

Foote, Figure S3

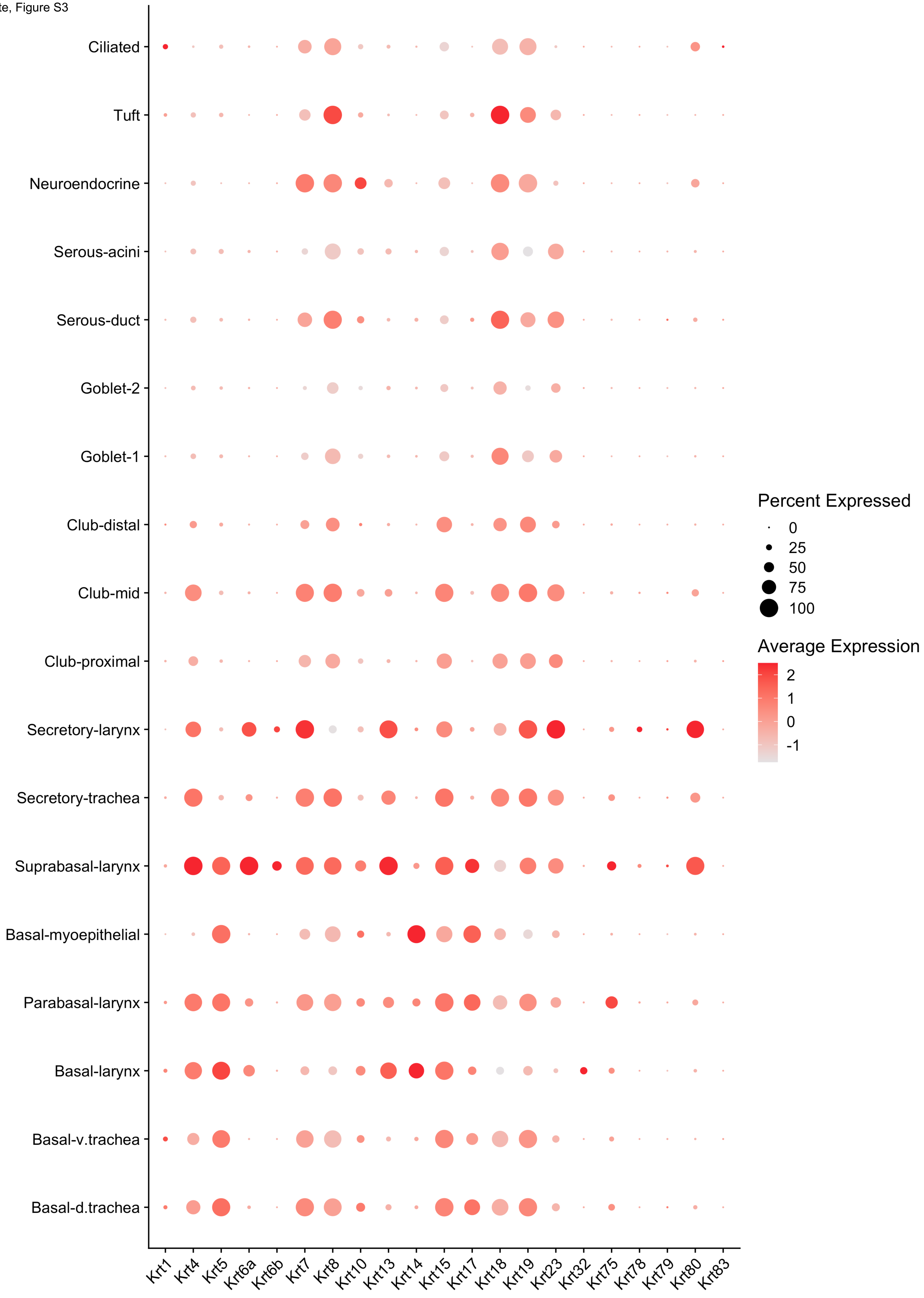

Foote, Figure S4

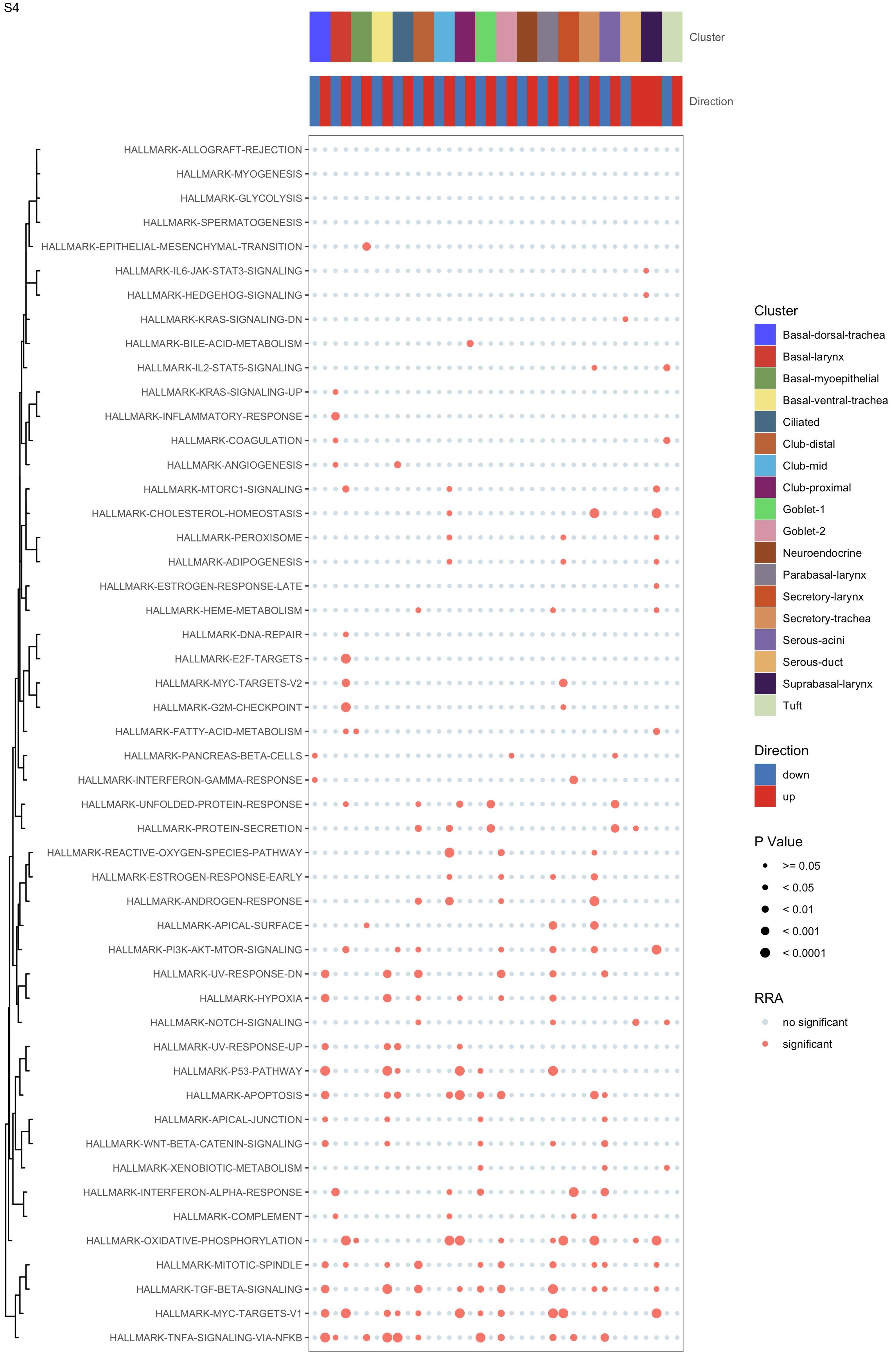

Foote, Figure S5

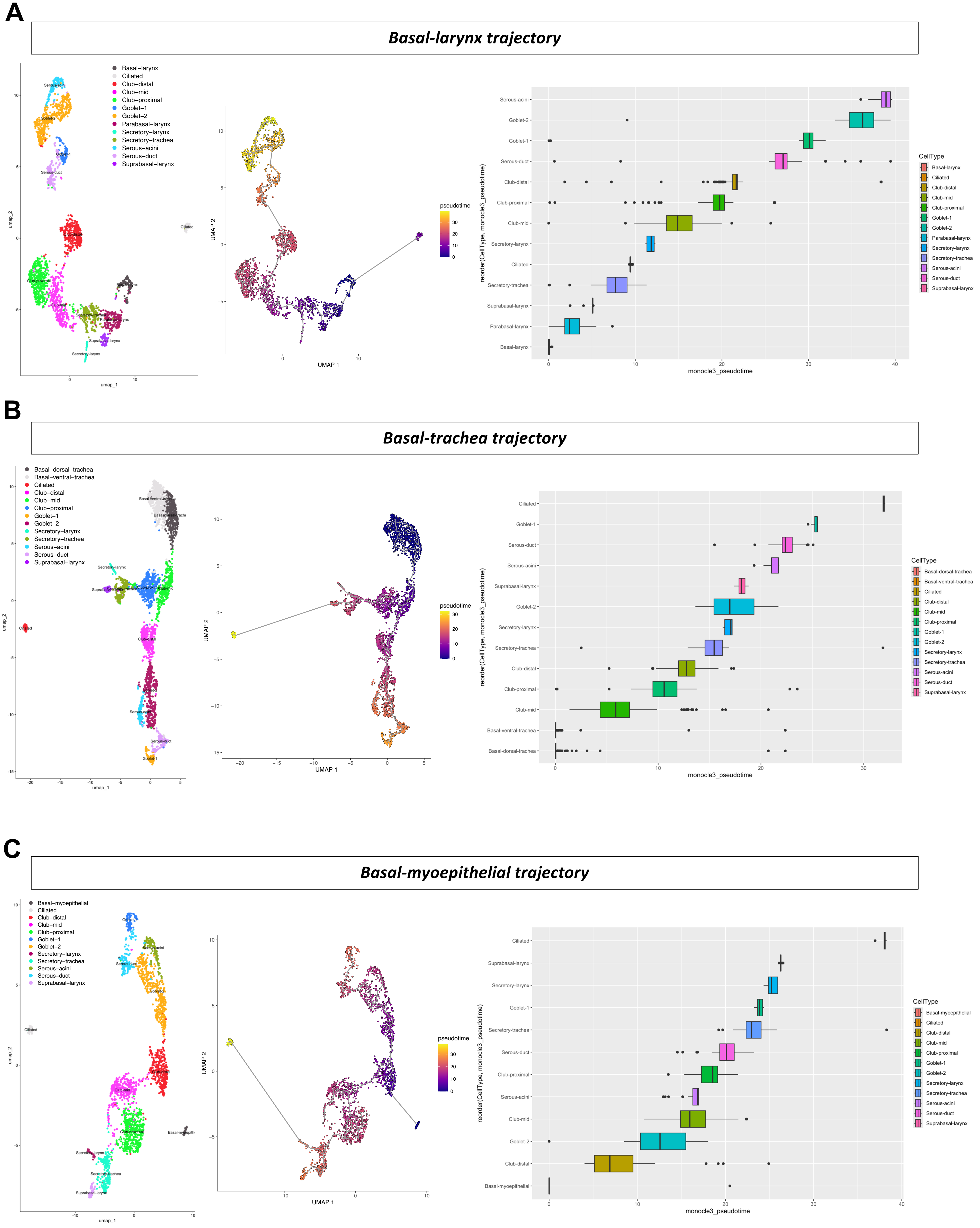
